# RNA editing controls gene drive by a Neurospora Spore killer

**DOI:** 10.1101/2020.12.30.424869

**Authors:** Shahriar Mahmud, Thomas M. Hammond, Nicholas A. Rhoades

**Affiliations:** School of Biological Sciences, Illinois State University, Normal, Illinois 61790

**Keywords:** fungi, gene driver, meiotic drive, RNA editing, spore killing

## Abstract

Neurospora *Sk-2* is a complex meiotic drive element that is transmitted to offspring through sexual reproduction in a biased manner. *Sk-2*’s transmission mechanism involves spore killing, and recent evidence has demonstrated that its spore killing mechanism is mediated by a gene called *rfk-1*. The native *rfk-1* sequence, referred to as *rfk-1^+^*, encodes an early UAG stop codon and a late UAA stop codon. When translation stops at the early stop codon, a 102 amino acid protein called RFK-1A is produced, and when translation stops at the late stop codon, a 130 amino acid protein called RFK-1B is produced. We show that expression of RFK-1B occurs when the early stop codon undergoes adenosine-to-inosine (A-to-I) mRNA editing (UAG is edited to UIG), and that this editing event is required for spore killing. We also show that RFK-1B, but not RFK-1A, acts as a poison when it is ectopically expressed within the vegetative tissue of a non-*Sk-2* strain. Furthermore, we show that RFK-1B toxicity can be neutralized in vegetative tissue by co-expressing RSK^Sk-2^, which is presumed to be *Sk-2*’s antidote protein. Finally, we show that *rfk-1^+^*’s first intron, or a truncated version of this intron, when present, improves phenotypic expression of RFK-1B in vegetative tissue. Overall, our results demonstrate that *Sk-2* uses A-to-I mRNA editing to control when its killer protein (poison) is produced, and that the primary killing and resistance functions of *Sk-2* can be established within a non-*Sk-2* strain by the ectopic expression of only two genes.

## INTRODUCTION

Neurospora *Spore killer-2* (*Sk-2*) is a 2.5 Mb selfish genetic element that achieves biased transmission to the next generation through spore killing (Raju 1979; Turner and Perkins 1979; Campbell and Turner 1987; Svedberg et al. 2018). Spore killing in Neurospora occurs during sexual development within fruiting bodies called perithecia. Each perithecium supports the development of hundreds of spore sacs, called asci, and each ascus supports the development of eight ascospores. In the absence of spore killing, mature asci typically contain eight viable black ascospores. However, when a cross is performed between an *Sk-2* strain and an *Sk-2*-sensitive strain (denoted as *Sk-S*), mature asci contain four viable black ascospores and four inviable hyaline ascospores, and the viable ascospores are almost always of the *Sk-2* genotype (>96%, Turner and Perkins 1979).

The Killer Neutralization (KN) model has been proposed to explain how *Sk-2* achieves biased transmission through spore killing (Hammond et al. 2012). This model holds that *Sk-2* expresses a killer protein (poison) and a resistance protein (antidote) throughout ascus development, up until ascospore delimitation, after which the antidote becomes restricted to *Sk-2* ascospores. The poison, however, is presumed to be either more stable than the antidote, or capable of export from *Sk-2* ascospores after delimitation. As a result, *Sk-2* ascospores survive because they possess the poison’s antidote, while S*k-S* ascospores are killed because they do not.

*Sk-2*’s antidote is presumed to be encoded by the *rsk* gene, and at least three classes of *rsk* alleles exist (Hammond et al. 2012). The *rsk^Sk-2^* class provides an antidote to *Sk-2’s* poison, while the *rsk^Sk-3^* class provides an antidote to the poison produced by *Spore killer-3,* a selfish genetic element that appears to share a common evolutionary origin with *Sk-2* (Turner and Perkins 1979; Svedberg et al. 2018). The third class of *rsk* alleles is found in *Sk-S* strains (*rsk^Sk-S^*), and while it remains possible that this class provides an antidote to an unidentified spore killer, mating and ascospore production is normal in *Sk-S rskΔ* homozygous crosses (Hammond et al. 2012).

Sequence analysis predicts that the *rsk^Sk-2^* allele of *Sk-2* strain FGSC 7426 encodes a 437 aa protein, while the *rsk^Sk-3^* allele in *Sk-3* strain FGSC 3194 encodes a 482 aa protein (Svedberg et al. 2018). The standard reference strain of *N. crassa*, FGSC 2489 (Galagan et al. 2003), is sensitive to both *Sk-2* and *Sk-3,* and thus its *rsk* allele is of the *rsk^Sk-S^* type. Sequence analysis of the *rsk^Sk-S^* allele from FGSC 2489 predicts that it encodes a 486 amino acid protein. All three classes of RSK proteins lack characterized protein domains, and how they function as antidotes remains unknown.

The *Sk-2* poison is encoded by a gene called *rfk-1.* This gene is located within a DNA interval called *AH36* at the right border of the *Sk-2* element (Figure 1, A and B; Harvey et al. 2014; Rhoades et al. 2019). The native *rfk-1* sequence (referred to as *rfk-1^+^*) encodes an early stop codon and a late stop codon (Figure 1B). Previous research has shown that the early stop codon (UAG) undergoes A-to-I mRNA editing to a tryptophan codon (UIG) in sexual tissue. However, a specific requirement of the edit for spore killing had been left uninvestigated. Apart from undergoing A-to-I mRNA editing, a second unusual feature of *rfk-1^+^* is its first intron, which contains seven repeats of a 46–48 nucleotide long sequence. It seems that these repeated sequences could influence *rfk-1*^+^ expression levels in a way that optimizes the ratio of poison to antidote during ascus development.

**Figure 1.**
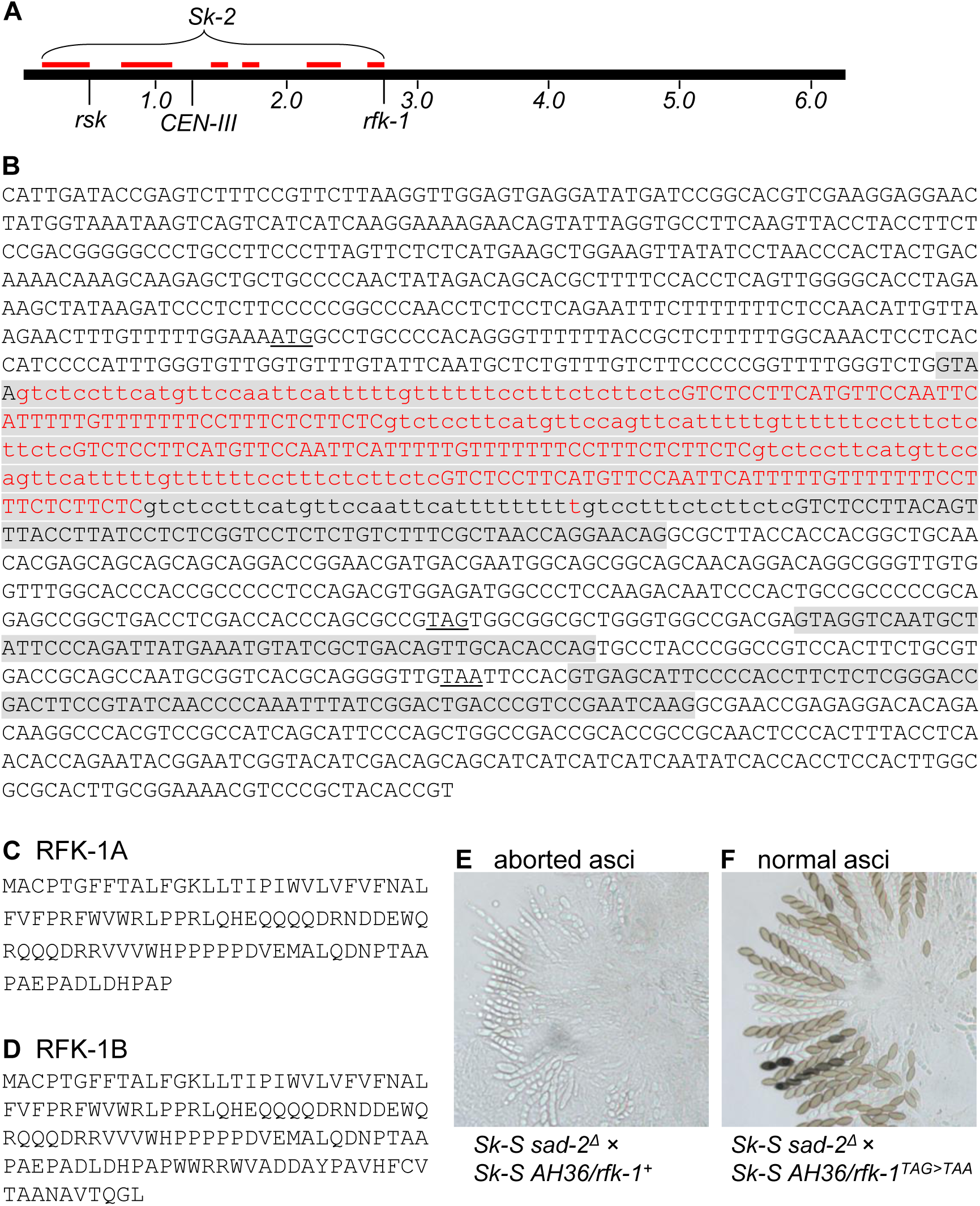
RNA editing of the early stop codon in *rfk-1^+^* transcripts to a tryptophan codon is required for spore killing. (A) A diagram of Chromosome III in *Sk-2* strains is shown. *Sk-2* spans approximately 2.5 Mb of Chromosome III. The antidote gene, *rsk*, is located on the left arm, while the poison gene, *rfk-1*, is located on the right arm. Inverted intervals within *Sk-2*, relative to Chromosome III in *Sk-S* strains, are indicated with red lines. These inversions are thought to suppress recombination within the *Sk-2* element. The diagram is based on a map of *Sk-2* provided by Svedberg et al. (2018). (B) The sequence of the *AH36* interval is shown (GenBank KJ908288.1: positions 27900–29380). The start and stop codons of *rfk-1^+^* are underlined. The three introns are indicated with gray shading. The first intron contains 7.2 repeats of a 46–48 bp sequence. The repeats are demarcated with alternating uppercase and lowercase letters. The bases deleted from the first intron in the *rfk-1* variant referred to as Int1T are marked with red font. (C) The predicted 102 amino acids of RFK-1A are shown. (D) The predicted 130 amino acids of RFK-1B are shown. (E) Asci from a cross between two *Sk-S* strains (ISU-3037 × ISU-4954), one of which carries the *AH36/rfk-1^+^* transgene, are shown. The presence of *AH36/rfk-1^+^* causes ascus abortion. The *sad-2Δ* allele was included in one of the crossing parents to suppress Meiotic Silencing by Unpaired DNA (Shiu et al. 2006). (F) Asci from a cross between two *SkS* strains (ISU-3037 × ISU-4955), one of which carries an *AH36/rfk-1^TAG>TAA^* transgene, are shown. Mutation of the early TAG stop codon to a TAA stop codon allows ascospores to develop.

In this study, we aimed to determine if A-to-I mRNA editing of the early *rfk-1^+^* stop codon is required for spore killing. We also aimed to identify the *Sk-2* poison and to determine if *Sk-2*’s poison could be phenotypically expressed in vegetative tissue. To complete these aims, we define two RFK-1 protein variants. The first, RFK-1A (Figure 1C), is produced when translation of *rfk-1^+^*transcripts is terminated at the early stop codon. The second, RFK-1B (Figure 1D), is produced when A-to-I mRNA editing of the early stop codon allows translation of *rfk-1^+^* transcripts to terminate at the late stop codon. By combining site-directed mutagenesis with spore killing assays, we show that A-to-I mRNA editing of the early stop codon, which allows for the production of RFK-1B, is required for spore killing. We also show that RFK-1B acts as a poison when expressed within the vegetative tissue of an *Sk-S* strain, and that RFK-1B can neutralized in vegetative tissue by expression of RSK^Sk-2^. Finally, we show that removal of *rfk-1*^+^’s first intron reduces phenotypic expression of RFK-1B in vegetative tissue, but that a truncated Intron 1, containing only one copy of the 46–48 nucleotide repeat, can be used in place of the native intron to obtain a level of RFK-1B expression that is at least equal to that obtained with the native intron if not more robust.

## MATERIALS AND METHODS

### Strains, media, and culture conditions

The genotypes of all strains used in this study are listed in Table 1. Vogel’s Minimal Medium (VMM) or Vogel’s Minimal Agar (VMA: VMM + 2% agar; Vogel 1956) was used for vegetative propagation of all fungal strains unless otherwise indicated. L-histidine was added to growth medium at 0.5 g per liter for culturing strains carrying a null *his-3* allele. Hygromycin B was used at 200 μg per ml when selecting for hygromycin resistance. Westergaard and Mitchell’s synthetic crossing agar (SCA; Westergaard and Mitchell 1947) with 1.5% sucrose and a pH of 6.5 was used for crosses. Brockman and de Serres agar (BDA; Brockman and De Serres 1963) with 1% sorbose, 0.05% glucose, 0.05% fructose, 1.5% agar, and the recommended salts/minerals, was used to induce colonial growth as needed. The *tcu-1* promoter was suppressed by adding an additional 250 μM of CuSO_4_ to culture medium. Below, because the *tcu-1* promoter was used to control expression of RFK proteins, VMA plus an additional 250 μM of CuSO_4_ is referred to as “RFK suppression” medium, and VMA without supplemental CuSO_4_ is referred to as “RFK expression” medium. All growth assays were performed at room temperature under ambient lighting.

**Table 1.**
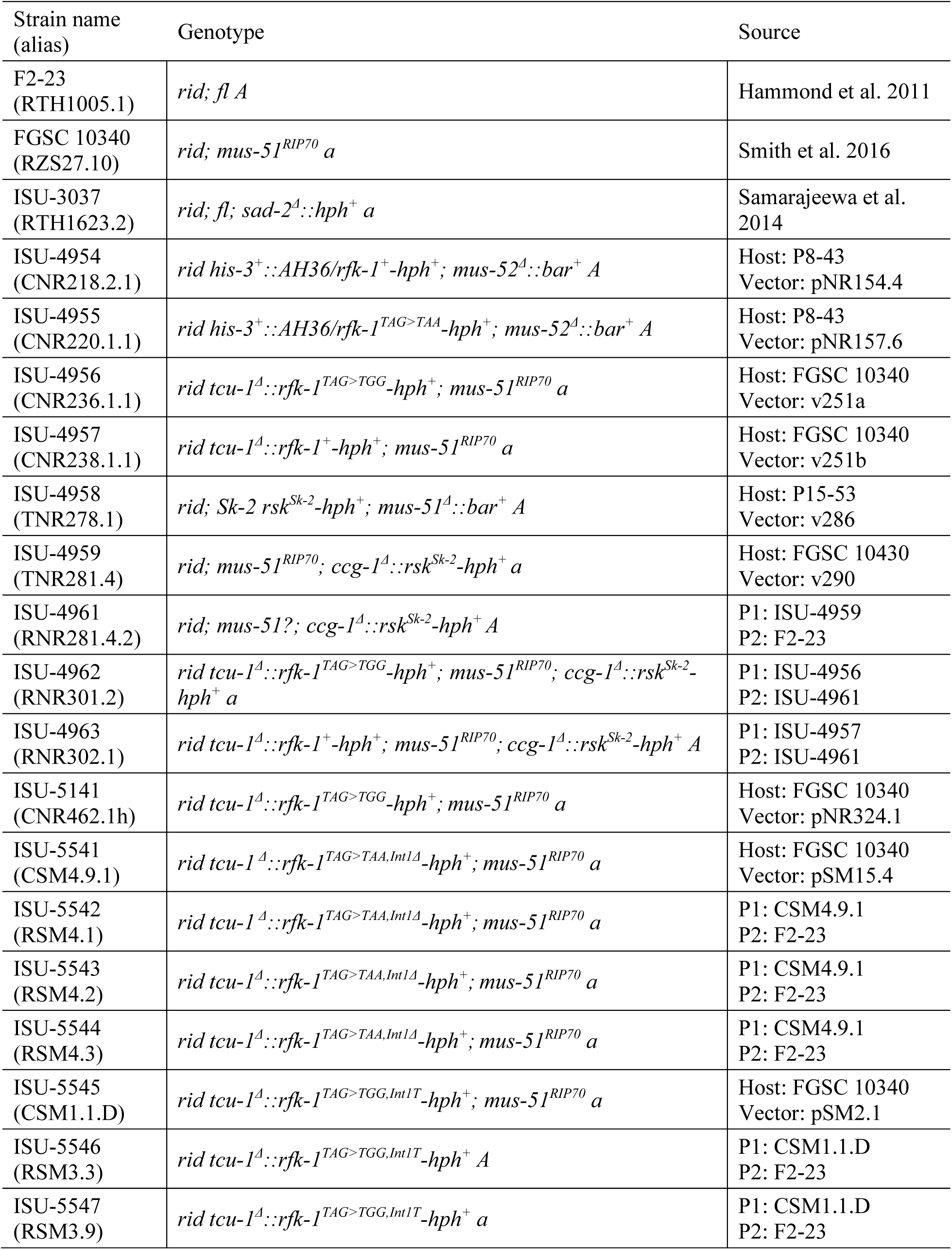
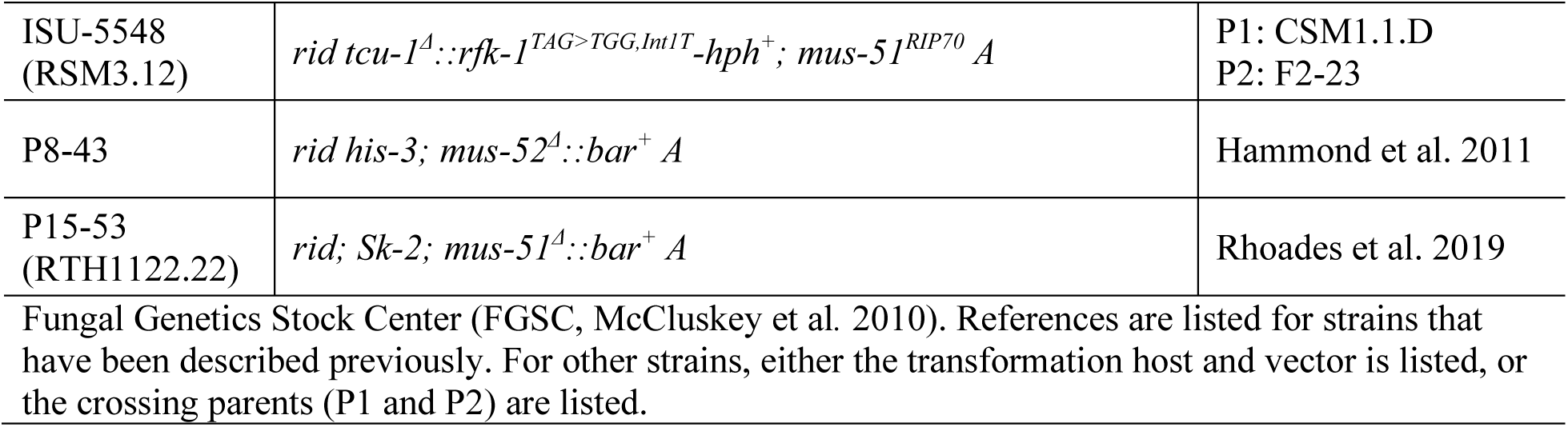
Strains used in this study.

### Plasmid construction

Sequences of PCR primers used in this study are listed in Table 2. Neurospora genome sequences were downloaded from FungiDB (Basenko et al. 2018) and/or the National Center for Biotechnology Information (NCBI; Sayers et al. 2025) unless otherwise indicated.

**Table 2.**
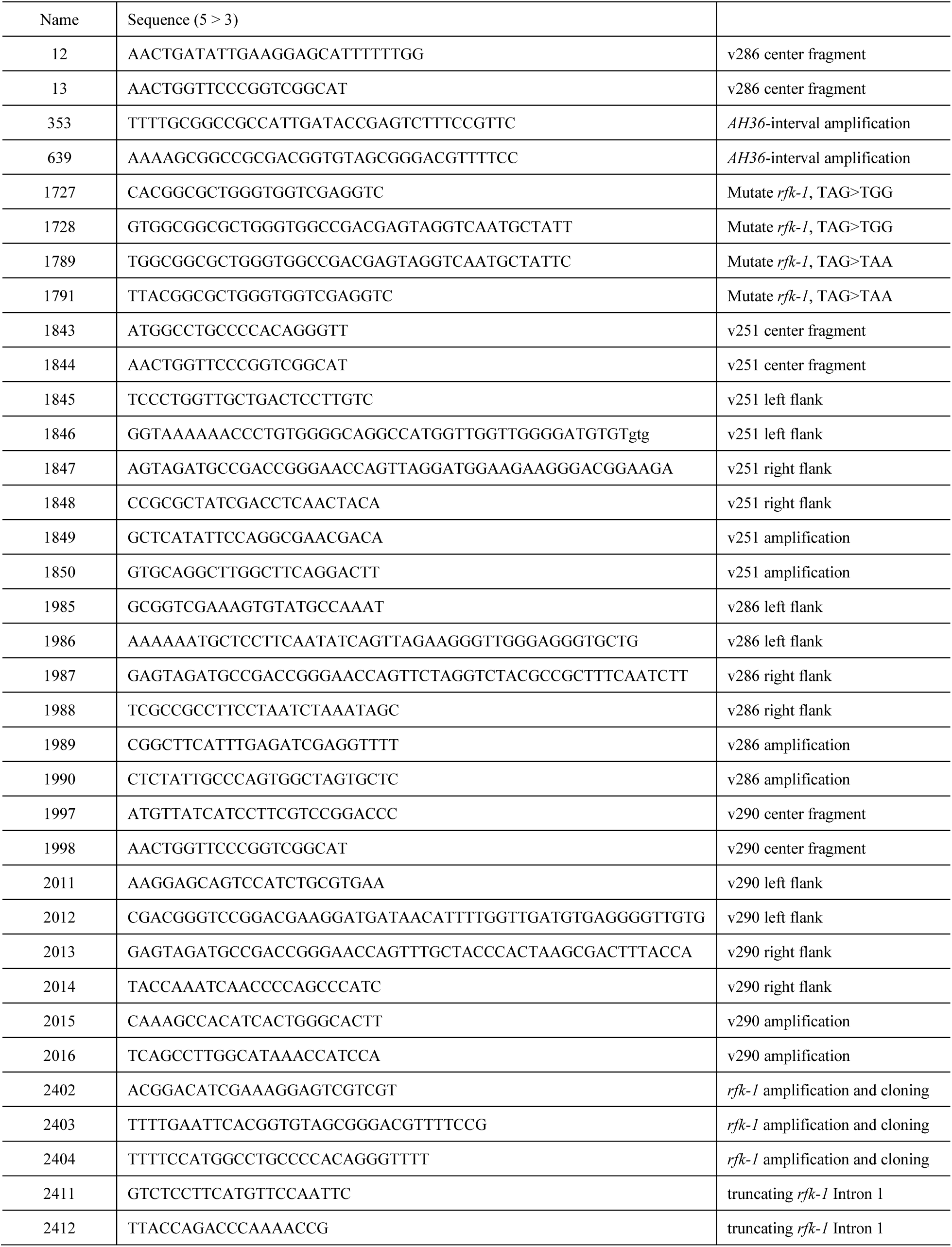
Primers used in this study.

#### Construction of plasmids for testing variants of the AH36 interval

Plasmid pNR9.1, which contains *AH36/rfk-1^+^* was constructed by PCR-amplifying the *AH36* interval from *Sk-2* genomic DNA with Primer Set 353/639, digesting the DNA molecule with *Not*I, and inserting the digested product into the *Not*I site of pBluescript II KS-(Stratagene). pNR127.9, which contains the *AH36/rfk-1^TAG>TGG^*, was constructed from pNR9.1 with Primer Set 1727/1728 and the Q5 Site-Directed Mutagenesis Kit (New England Biolabs). Similarly, pNR153.1, which contains *AH36/rfk-1^TAG>TAA^*, was constructed from pNR9.1 with Primer Set 1789/1791 and the Q5 Site-Directed Mutagenesis Kit. pNR154.4 and pNR154.6 were constructed by transferring the *Not*I fragment containing *AH36*/*rfk-1^+^* from pNR9.1 to the *Not*I site of pTH1256.1 (GenBank, MH550659.1). The *rfk-1^+^* and *hph^+^* coding sequences are in opposite orientations in pNR154.4 and identical orientations in pNR154.6. pNR155.2 was constructed by transferring a 1.5 kb *Not*I fragment containing *AH36/rfk-1^TAG>TGG^* from pNR127.9 to the *Not*I site of pTH1256.1. The *rfk-1^TAG>TGG^* and *hph^+^*coding sequences are in identical orientations in pNR155.2. pNR157.6 was constructed by transferring a 1.5 kb *Not*I fragment containing *AH36/rfk-1^TAG>TAA^* from pNR153.1 to the *Not*I site of pTH1256.1. The *rfk-1^TAG>TAA^* and *hph^+^* coding sequences are in opposite orientations in pNR157.6.

#### Construction of plasmids for fusing the tcu-1 promoter to rfk-1 variants

pNR282.1 was constructed by inserting Synthetic DNA 1 (Figure S1, Integrated DNA Technologies) into pJET1.2 with the CloneJET PCR Cloning Kit. Synthetic DNA 1 contains *AH36/rfk-1^+,Int1Δ^.* pSM3.8 contains *AH36/rfk-1^TAG>TGG,Int1Δ^.* It was constructed from pNR282.1 with Primer Set 1727/1728 and the Q5 Site-Directed Mutagenesis Kit. pSM12.1 was constructed by inserting a PCR product obtained from pSM3.8 with Primer Set 2403/2404 into pJET1.2 with the CloneJET PCR Cloning Kit. pSM15.4 was constructed by digesting pSM12.1 with *Nco*I and *Eco*RI, and inserting the released DNA molecule containing *rfk-1^TAG>TGG,Int1Δ^*between the *Nco*I and *Eco*RI sites of a *tcu-1* replacement vector called pNR304.1 (GenBank PX833383). pNR321.1 was constructed by inserting the PCR product obtained from genomic DNA of ISU-4956 with Primer Set 2402/2403 into pJET1.2 with the CloneJET PCR Cloning Kit. pSM1.9 contains *rfk-1^TAG>TGG,Int1T^*. It was generated from pNR321.1 with Primer Set 2411/2412 and the Q5 Site-Directed Mutagenesis Kit. pSM2.1 was constructed by digesting pSM1.9 with *Nco*I and *Eco*RI and inserting the released DNA molecule containing *rfk-1^TAG>TGG,Int1T^* between the *Nco*I and *Eco*RI sites of pNR304.1. Similarly, pNR324.1 was constructed by amplifying a PCR product from genomic DNA of ISU-4956 with Primer Set 2402/2403, digesting the product with *Nco*I and *Eco*RI, and cloning the digested product between the *Nco*I and *Eco*RI recognition sites of pNR304.1.

### Double joint fusion PCR-based construction of transformation vectors

Double joint fusion PCR (DJ-PCR) was performed with the method of Yu et al. (2004). Transformation Vector v251a was designed to fuse the coding region of *rfk-1^TAG>TGG^* to the promoter of the *tcu-1* gene on Chromosome I of *Sk^S^* strain FGSC 10340 while deleting *tcu-1* coding sequences (Figure S2A). The left and right flanks of v251a were PCR-amplified from FGSC 10340 genomic DNA with Primer Sets 1845/1846 and 1847/1848, respectively. The center fragment was amplified from pNR155.2 with Primer Set 1843/1844. The flanks were fused to the center fragment, and the resulting fusion product was PCR-amplified with Primer Set 1849/1850. Transformation Vector v251b was constructed with the same method used to construct v251a, except that the center fragment was PCR-amplified from pNR154.6 instead of pNR155.2. Transformation Vector v286 was designed to insert *hph^+^* downstream of the *rsk* gene on Chromosome III in an *Sk-2* strain (Figure S2B). The left and right flanks were amplified from genomic DNA of P15-53 with Primer Sets 1985/1986 and 1987/1988, respectively. The center fragment was amplified from pTH1256.1 with Primer Set 12/13. The flanks were fused to the center fragment, and the resulting fusion product was PCR-amplified with Primer Set 1989/1990. Transformation Vector v290 was designed to fuse the coding region of *rfk-1^+^*to the *ccg-1* promoter on Chromosome V of an *Sk-S* strain while deleting the *ccg-1* coding sequence (Figure S2C). The left and right flanks were amplified from FGSC 10340 genomic DNA with Primer Sets 2011/2012 and 2013/2014, respectively. The center fragment was amplified from genomic DNA of ISU-4958 with Primer Set 1997/1998. The flanks were fused to the center fragment, and the resulting fusion product was PCR-amplified with Primer Set 2015/2016.

### *N. crassa* transformations

Transformation of *N. crassa* was performed with the electroporation method of Margolin et al. (1997) with modifications as described by Rhoades et al. (2020). Homokaryotic transformants were obtained by purification of heterokaryons through microconidia (Ebbole and Sachs 1990), macroconidia, or ascospores, and homokaryotic genotypes were confirmed by DNA isolation and PCR-based genotyping. ISU-4954 and ISU-4955 were obtained by transforming P8-43 with pNR154.4 and pNR157.6, respectively. ISU-4958 was obtained by transforming P15-53 with v286. The following strains were obtained by transformation of FGSC 10340: ISU-4956 (with v251a), ISU-4957 (with v251b), ISU-4959 (with v290), ISU-5141 (with pNR324.1), ISU-5541 (with pSM15.4), and ISU-5545 (with pSM2.1).

### Growth assays

Conidial suspensions were made in sterile water from cultures that were started at the same time on the same batch of RFK Suppression Medium. Conidial suspensions of 200 conidia/μl and 1000 conidia/μl were made for radial growth assays and linear growth assays, respectively.

Radial growth assays were performed by inoculating the centers of 100 mm culture dishes containing 25.0 ml of RFK Expression Medium or RFK Suppression Medium with 5 μl of conidial suspension (1000 conidia total). Inoculated plates were imaged after 96–120 hours of growth. Linear growth assays were performed in 25 ml polystyrene serological pipettes containing RFK Expression Medium or RFK Suppression Medium according to the method of White and Woodward (1995). The pipettes were inoculated with 5 μl of conidial suspension (5000 conidia total) at one end and the leading edges of the cultures were recorded at 12-hour intervals.

### Spore killing assays

Unidirectional crosses were performed on SCA as previously described (Samarajeewa et al. 2014). Asci were dissected from fruiting bodies into 50% glycerol on the 14^th^ day after fertilization. Digital images of asci were taken with a Leica DMBRE microscope and imaging system.

## RESULTS

### RFK-1B is required for spore killing

*rfk-1*’s discovery was made possible by first refining its location to a 1481 bp interval called *AH36* (Figure 1B). This interval contains *rfk-1* promoter sequences, *rfk-1*’s coding region, and *rfk-1* terminator sequences. Interestingly, its insertion into the genome of an *Sk-S* strain has no obvious effect on vegetative growth or developmental processes (Rhoades et al. 2019). However, when an *AH36*-harboring *Sk-S* strain is mated with another *Sk-S* strain, the cross produces asci with an aborted phenotype (Figure 1E; Rhoades et al. 2019). This suggests that the *AH36* transgene directs the production of the *Sk-2* poison during ascus development, but not at other stages of the Neurospora life cycle.

To determine if the poison produced by the *AH36* transgene requires the early stop codon within *rfk-1^+^* transcripts to undergo A-to-I mRNA editing, we used site directed mutagenesis to change the early stop codon within *rfk-1^+^* from TAG to TAA. Because the second position of the early stop codon is the one that undergoes A-to-I mRNA editing (i.e., UAG becomes UIG; where I is interpreted as G by the translational machinery; Rhoades et al. 2019), we reasoned that changing the third position to A would cause the early stop codon to remain a stop codon even if its second position undergoes editing (i.e., UAA becomes UIA). Thus, transcripts from an *AH36/rfk-1^TAG>TAA^* transgene should only be capable of directing the production of RFK-1A. When we examined a cross between a transformant carrying the *Sk-S AH36/rfk-1^TAG>TAA^*transgene and an *Sk-S* strain, we found that asci developed normally (Figure 1F). This finding demonstrates that the ascus abortion phenotype of *Sk-S AH36/rfk-1^+^* × *Sk-S* crosses (Figure 1E) requires the early stop codon in *rfk-1^+^* transcripts to undergo editing in a way that allows translational readthrough to the late stop codon. This finding also demonstrates that RFK-1A is insufficient for inducing ascus abortion.

We next sought to determine if RFK-1B can induce ascus abortion. To do so, we constructed an *AH36/rfk-1^TAG>TGG^* transgene, in which the early *rfk-1^+^* stop codon has been changed to a tryptophan-coding codon. We reasoned that this transgene should only be capable of directing the production of RFK-1B. However, our attempts to transform an *Sk-S* strain with this *AH36/rfk-1^TAG>TGG^* transgene failed to produce transformants, possibly because RFK-1B is toxic and *AH36/rfk-1^TAG>TGG^*directs the production of RFK-1B in transformed tissue.

### RFK-1B is toxic to Neurospora vegetative tissue

To specifically test if either RFK-1A or RFK-1B is toxic to vegetative tissue, we placed *rfk-1^+^* and *rfk-1^TAG>TGG^*coding sequences under the control of the copper repressible *tcu-1* promoter (Lamb et al. 2013) in an *Sk-S* genetic background (Figure S2A). When driven by the *tcu-1* promoter, *rfk-1^+^* transcripts should only direct the production of RFK-1A because A-to-I mRNA editing does not occur outside of the sexual phase (Liu et al. 2017). Furthermore, *rfk-1^TAG>TGG^*transcripts should only direct the production of RFK-1B because the early stop codon has been changed to a tryptophan codon. When examined in growth assays, we found that the RFK-1A strain (*tcu-1^Δ^::rfk-1^+^*) grew similarly to the control strain (*tcu-1+*) under RFK suppression and expression conditions (Figure 2, A and B, ISU-4957). In contrast, the RFK-1B strain (*tcu-1^Δ^::rfk-1^TAG>TGG^)* grew similarly to the control strain only under RFK suppression conditions. Under RFK expression conditions, growth of the RFK-1B strain was considerably restricted (Figure 2, ISU 4956). These results demonstrate that RFK-1B expression is toxic to vegetative tissue.

**Figure 2.**
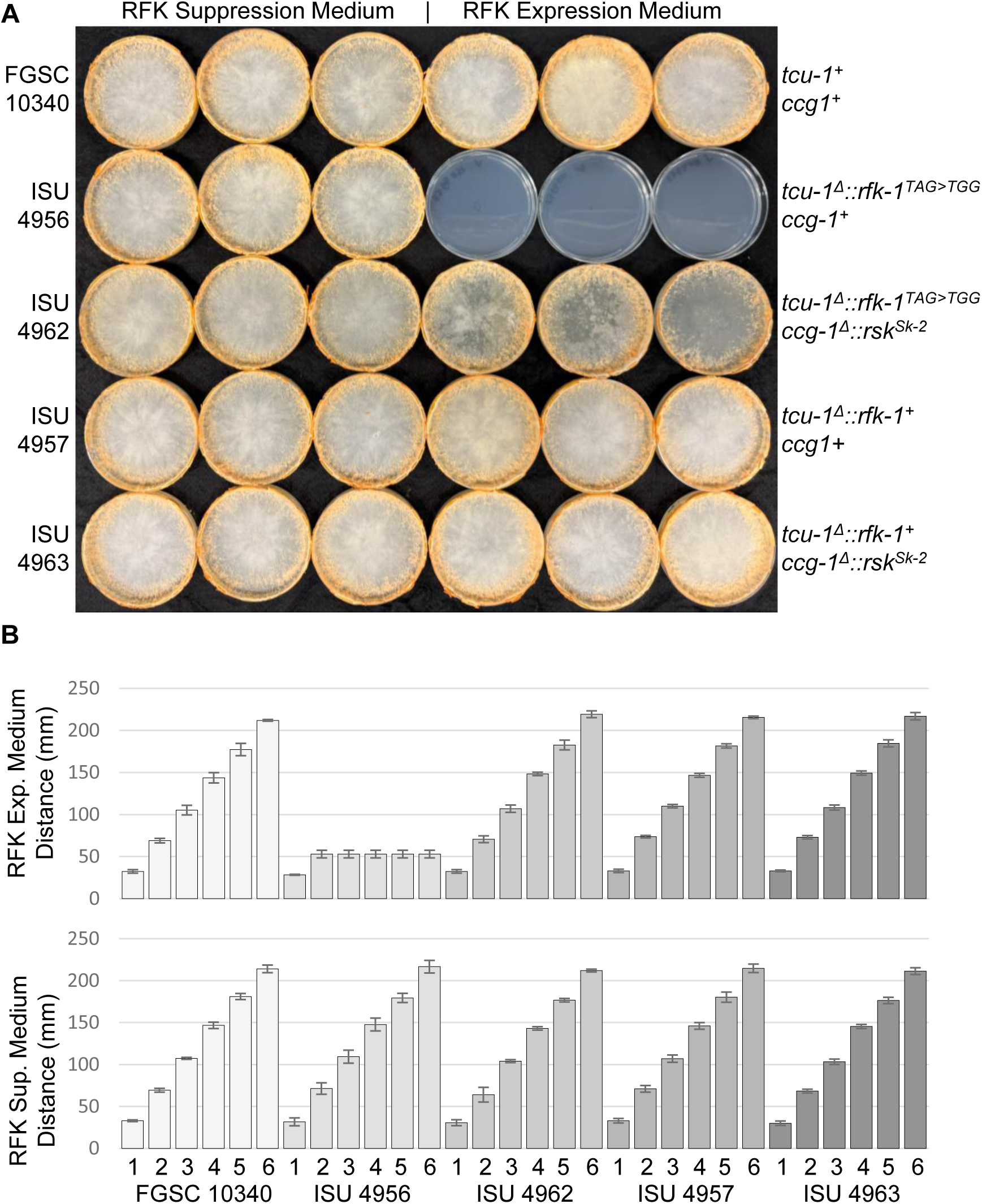
Phenotypic expression of RFK-1B and RSK^Sk-2^ in vegetative tissue. (A) Radial growth assays were performed on RFK Suppression Medium (high copper) and RFK Expression Medium (low copper). Three technical replicates were performed for each strain under each culture condition. The image was taken five days after point inoculation at the center of each culture dish. Strain names and key genotypes are indicated to the left and right of the image, respectively. The *tcu-1^Δ^* alleles were created by replacing *tcu-1* coding sequence with *rfk-1* coding sequences, thereby placing the *rfk-1* coding sequences under control of the copper repressible *tcu-1* promoter. The *ccg-1^Δ^* allele was created by replacing the *ccg-1* coding sequence with the *rsk^Sk-2^*coding sequence, thereby placing the *rsk^Sk-2^* coding sequence under control of the *ccg-1* promoter. These results indicate that *tcu-1^Δ^::rfk-1^TAG>TGG^* strains grow poorly on RFK Expression Medium (when RFK-1B is expressed), unless the strain also carries *ccg-1^Δ^::rsk^Sk-2^*(which expresses RSK^Sk-2^, presumably in both types of medium). In contrast, the growth of *tcu-1^Δ^::rfk-1^+^* strains is similar on both types of medium and unaffected by the presence of *ccg-1^Δ^::rsk^Sk-2^*. (B) Linear growth assays were performed with the same strains depicted in Panel A. Bars represent distances traveled at the 48-, 60-, 72-, 84-, 96-, and 108-hour time points. Distances were normalized to the position of each culture’s growth front at 36 hours. Three replicates were performed for each strain. Error bars are standard deviation. The results are consistent with the radial growth assay results presented in Panel A.

### RSK^Sk-2^ neutralizes RFK-1B in vegetative tissue

RFK-1B’s toxicity, along with RFK-1A’s apparent innocuousness, suggests that RFK-1B is *Sk-2*’s poison. If true, then concomitant expression of *Sk-2*’s antidote, RSK^Sk-2^, in vegetative tissue should neutralize RFK-1B. To test this possibility, we placed the coding region for *rsk^Sk-2^* under control of the *ccg-1* promoter in an *Sk-S* genetic background (Figure S2C). This promoter is routinely used for obtaining robust expression of transgenes in *N. crassa* (McNally and Free 1988; Loros et al. 1989; Bell-Pedersen et al. 1996). We then crossed the resulting *rsk^Sk-2^*strain with an RFK-1A-expressing strain and an RFK-1B-expressing strain to isolate offspring capable of expressing either RFK-1A and RSK^Sk-2^, or RFK-1B and RSK^Sk-2^. As expected, the RFK-1A offspring that carries *rsk^Sk-2^* grew similarly under both RFK suppression and expression conditions (Figure 2, ISU-4963), just like the RFK-1A strain that lacks *rsk^Sk-2^*(Figure 2, ISU-4957). Interestingly, the RFK-1B offspring that carries *rsk^Sk-2^*also grew similarly under RFK suppression and expression conditions (Figure 2, ISU-4962). This is in stark contrast to the RFK-1B strain that lacks *rsk^Sk-2^*, whose growth was restricted under RFK-1B expression conditions (Figure 2, ISU-4956). Together, these results demonstrate that RFK-1B expression is vegetatively toxic and that this toxicity can be neutralized by RSK^Sk-2^.

### The first intron of *rfk-1^+^* influences phenotypic expression of RFK-1B in vegetative tissue

Our initial procedure for placing *rfk-1* variants under the control of *tcu-1(P)*, described above, involved constructing transformation vectors with a technique called DJ-PCR (Yu et al. 2004). To simplify the process of testing *rfk-1* variants, we created pNR304.1 (GenBank: PX833383; Figure S3). This plasmid can be used to construct transformation vectors that replace the *tcu-1* coding sequence on Chromosome I with the coding sequence of any gene. As described below, we used pNR304.1 to examine the role of *rfk-1^+^*’s first intron in vegetative expression of RFK-1B.

One of the most unusual aspects of the *rfk-1^+^* coding region is its first intron, which contains 7.2 copies of a 46–48 bp long repeat (Figure 1B). The unusual structure of this intron hints that it may be important for the success of gene drive by spore killing. We sought to test the importance of the first intron by placing three *rfk-1* variants under control of the *tcu-1* promoter. The first variant, which served as the positive control for vegetative expression of RFK-1B, contains a full-length native Intron 1 and is referred to as Int1+. This *rfk-1* variant is essentially identical to the *rfk-1^TAG>TGG^*variant in Figure 2 (e.g., ISU-4956) except that it was constructed with pNR304.1 rather than DJ-PCR. The second variant lacks Intron 1 and is referred to as Int1Δ. The third variant contains only one of the 46–48 bp repeats and is referred to as Int1T (Figure 1A: the red font within Intron 1 marks the deleted bases in Int1T). The three variants were then compared in growth assays. As expected from our previous tests of *rfk-1^TAG>TGG^*, growth of the Int1+ strain was restricted under RFK expression conditions (Figure 3, ISU-5141). However, and somewhat surprisingly, Int1Δ’s growth was less restricted than Int1+ under the same conditions. For example, the results presented in Figure 3 compare the growth of an Int1+ strain to three Int1Δ strains (three siblings from the same cross). Each Int1Δ strain formed more tissue than the Int1+ strain on RFK Expression Medium in the radial growth assay (Figure 3A), and each Int1Δ strain covered more distance than the Int1+ strain on RFK Expression Medium in the linear growth assay (Figure 3B). These results indicate that the presence of Intron 1 improves phenotypic expression of RFK-1B in vegetative tissue. We therefore designed the Int1T variant to examine how removing all but one of the repeats within Intron 1 would influence phenotypic expression of RFK-1B. We found that each of the three tested Int1T strains (siblings from the same cross) grew more poorly than the Int1+ control under RFK expression conditions. For example, in the radial growth assay, mycelia is present across the surface of the RFK Expression Medium for the Int1+ strain, but not for any of the Int1T strains on the same medium (Figure 4A, the sparse mycelia across the surfaces of the RFK Expression Medium for the Int1+ replicates may be difficult to see in the image). Additionally, each of the Int1T strains covered less distance on RFK Expression Medium than the Int1+ strain in the linear growth assay (Figure 4B).

**Figure 3.**
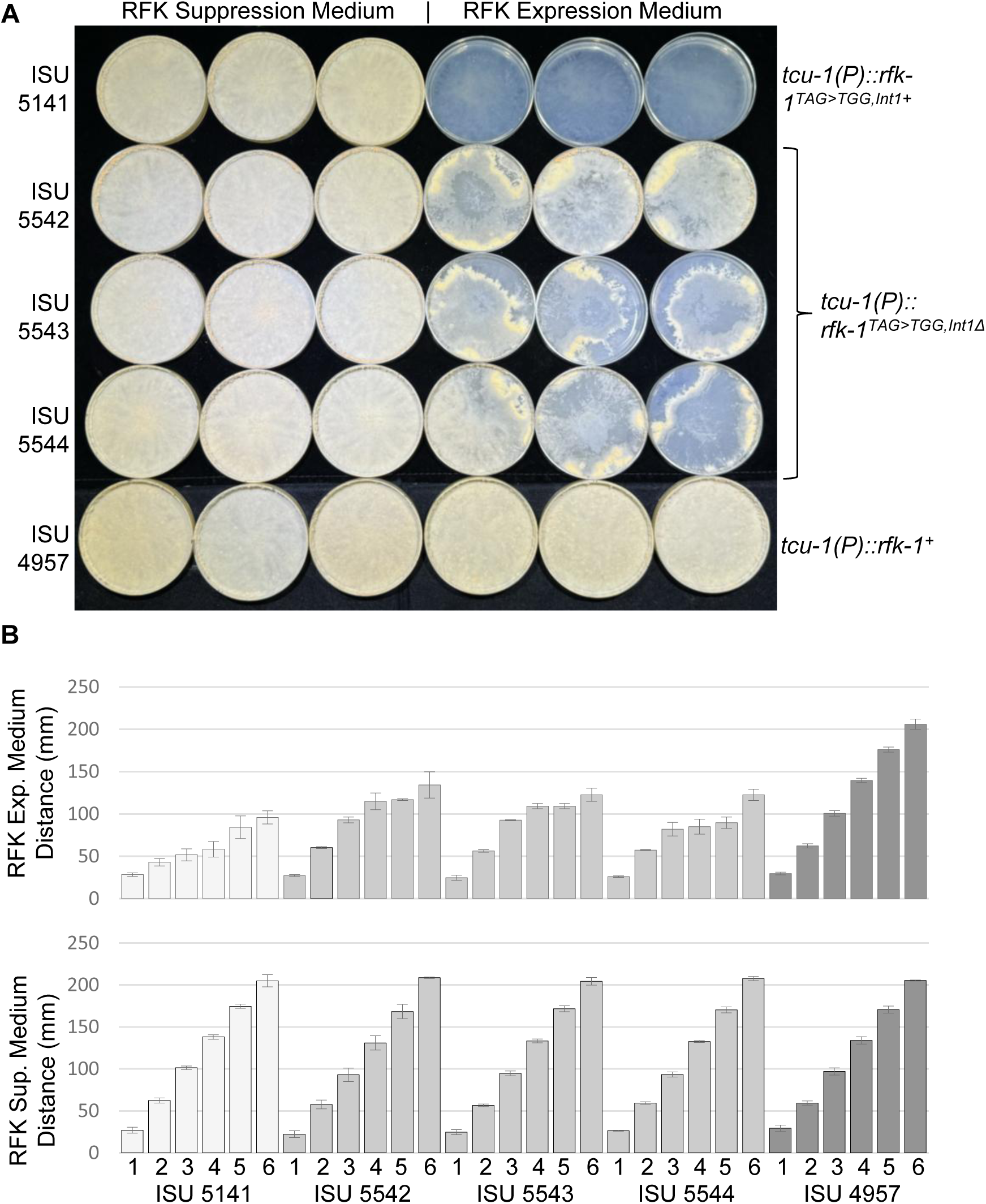
Deletion of *rfk-1^+^’s* first intron reduces phenotypic expression of RFK-1B in vegetative tissue. (A) Radial growth assays were performed as described in Figure 2. ISU-5542, ISU-5543, and ISU-5544 are siblings. These results show that *tcu-1^Δ^*::*rfk-1^TAG>TGG,Int1+^* produces a more robust RFK-1B expression phenotype than does *tcu-1^Δ^*::*rfk-1^TAG>TGG^*^,*Int1Δ*^. (B) Linear growth assays were performed for the strains depicted in Panel A as described in Figure 2. These results are consistent with the radial growth assay results presented in Panel A.

**Figure 4.**
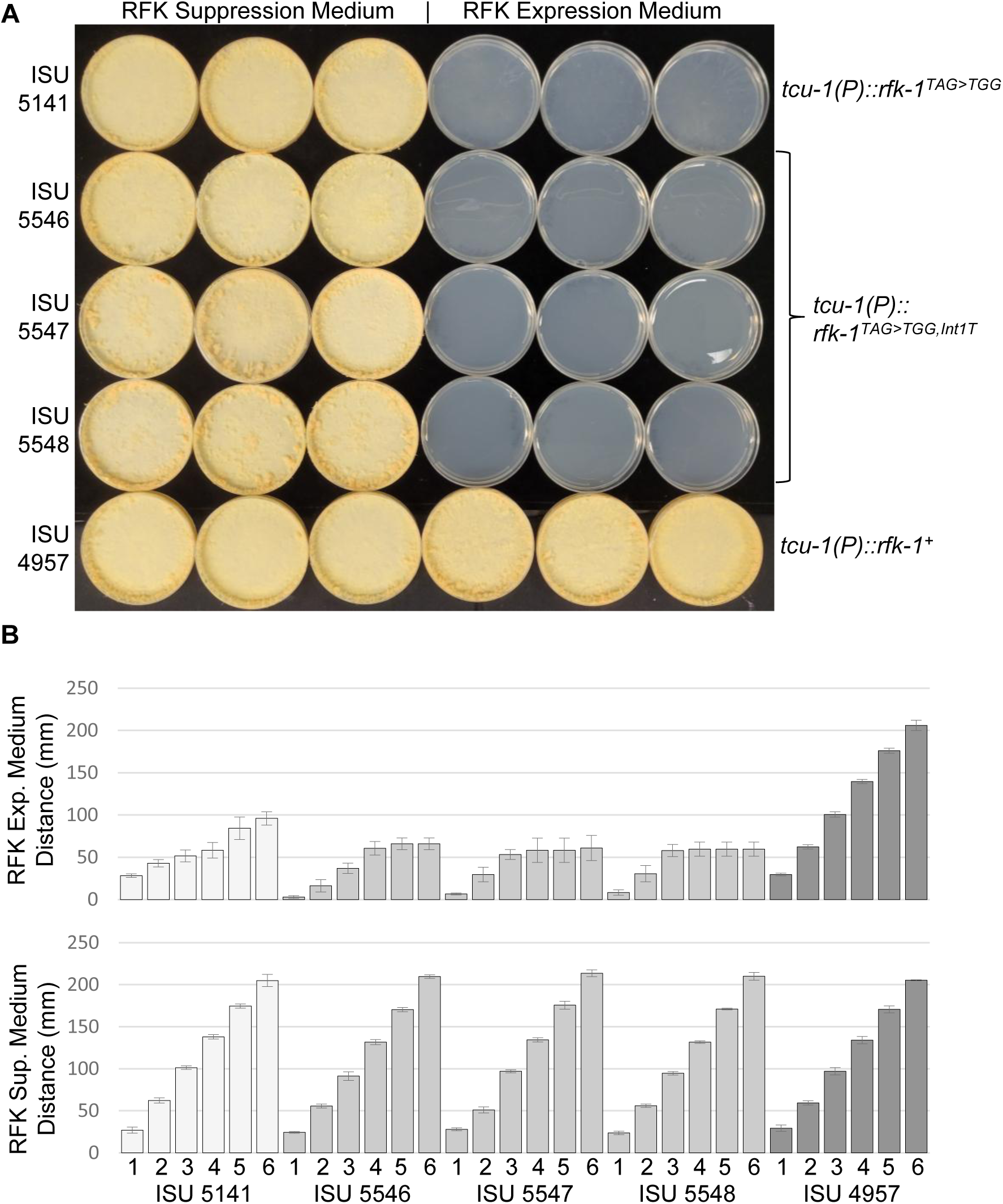
A truncated *rfk-1* intron provides robust phenotypic expression of RFK-1B in vegetative tissue. (A) Radial growth assays were performed as described in Figure 2. ISU-5546, ISU-5547, and ISU-5548 are siblings. These results show that *tcu-1^Δ^*::*rfk-1^TAG>TGG^*^,*Int1T*^ provides for robust phenotypic expression of RFK-1B at a level that is at least equal to, if not greater than, *tcu-1^Δ^*::*rfk-1^TAG>TGG,Int1+^*on RFK Expression Medium. While it may be difficult to detect in the image, sparse mycelia is present over the surface of each *rfk-1^TAG>TGG,Int1+^*replicate but not each *rfk-1^TAG>TGG^*^,*Int1T*^ replicate. (B) Linear growth assays were performed for the strains depicted in Panel A as described in Figure 2. These results are consistent with the radial growth assay results presented in Panel A.

Overall, these results demonstrate that an *rfk-1^TAG>TGG^* variant containing either Int1+ or Int1T phenotypically expresses RFK-1B in vegetative tissue more robustly than an *rfk-1^TAG>TGG^* variant lacking Intron 1.

## DISCUSSION

Killer meiotic drivers (KMDs) are widespread in ascomycete fungi. They achieve drive by killing spores that fail to inherit the driving element. Well known examples include *het-s* and the Spok family in Podospora fungi, *tdk1* and the *wtf* family in Schizosaccharomyces fungi, and the spore killers in Neurospora fungi (Zanders and Johannesson 2021). The mechanisms by which the known KMDs achieve drive appear highly diverse. The *het-s* mechanism involves prion formation (Dalstra et al. 2003), the *tdk1* mechanism involves disrupting mitosis (Hua, Zhang, Yang, Zhang, et al. 2024; Hua, Zhang, Yang, Ren, et al. 2024), the *wtf* drivers form toxic aggregates (Eickbush et al. 2019; Nuckolls et al. 2020; Nunez et al. 2020), and the Spok family members degrade DNA (van der Gaag et al. 2000; Vogan et al. 2019; Urquhart and Gardiner 2023). The mechanisms used by the Neurospora Spore killers are perhaps the least understood.

In this study, we have provided evidence that *Sk-2*’s killer protein (poison) is RFK-1B and that expression of RFK-1B is controlled by A-to-I mRNA editing. Furthermore, we found that RFK-1B and RSK^Sk-2^, *Sk-2*’s antidote, can function in vegetative tissue. Our results also demonstrate that phenotypic vegetative expression of RFK-1B is more robust when *rfk-1^+^’s* native first intron, or a truncated version of this intron, is present in *rfk-1* pre-mRNAs than when it is completely removed.

A-to-I mRNA editing has been identified in several Pezizomycotinia fungi in addition to *N. crassa*, including *Fusarium graminearum*, *Fusarium verticillioides*, *Sordaria macrospora*, and *Pyronema confluens* (Liu et al. 2016; Liu et al. 2017; Teichert et al. 2017). Like alternate splicing, A-to-I mRNA editing is thought to be beneficial because it can provide an ability to encode multiple, useful proteins from a single gene. Major support for this hypothesis is found in *F. graminearum,* a fungus that requires optimal expression of both unedited and edited transcripts of a gene called *CME11* to maximize reproductive success (Xin et al. 2023).

However, A-to-I mRNA editing is not conserved across the fungal kingdom. The reason may be that there are fitness disadvantages associated with the mechanism. For example, our results here demonstrate that *Sk-2* has coopted the A-to-I mRNA editing machinery to control where and when its poison is produced. This is the second example of a fungal spore killer that has taken advantage of its host’s A-to-I mRNA editing machinery to break the rules of Mendelian inheritance. The other known example is *Skc1*, an *F. verticillioides* spore killer that requires A-to-I mRNA editing of an early stop codon to express the toxic version of its killer protein (Lohmar et al. 2022). *Skc1* is a single gene spore killer, and how it kills spores while resisting killing is unknown.

Unlike *Skc1*, *Sk-2* requires multiple gene products to be transmitted in a biased manner to offspring through spore killing. While the KN model predicts that the *Sk-2* antidote (RSK^Sk-2^) neutralizes the *Sk-2* poison (RFK-1B) through a direct interaction, a direct interaction between these two proteins has yet to be demonstrated. Still, the results presented here suggest that RFK-1B neutralization requires only RSK^Sk-2^ because neutralization of RFK-1B in an *Sk-S* background only requires ectopically expressing RSK^Sk-2^ in the same genetic background.

However, given that *Sk-2* is a complex meiotic drive element that spans hundreds of protein-coding genes (Svedberg et al. 2018), it remains possible that some of these genes modify the efficiency of RFK-1B-mediated killing and/or RSK^Sk-2^-mediated resistance. Testing this hypothesis will likely require construction of a functional two gene drive element containing only *rfk* and *rsk* genes, then comparing the ability of this minimal element to drive with that of the native *Sk-2* element.

The *rfk-1* gene appears to have evolved from a partial duplication of gene *ncu07086*, resulting in the first 39 amino acids of both RFK-1A and RFK-1B being similar to the first 39 amino acids of NCU07086 (Rhoades et al. 2019). Although NCU07086 has low homology to proteins of the AtpF Superfamily, its specific function in Neurospora has yet to be elucidated. The first intron of *rfk-1*^+^ also appears to have originated from *ncu07086*. For example, all but six nucleotides of the 46–48 bp repeat with *rfk-1* Intron 1 are identical to a 47 bp sequence within *ncu07086* first intron (Rhoades et al. 2019). While we are unsure why phenotypic expression of RFK-1B in vegetative tissue is more robust when Intron 1 is present than when it is removed, our identification of a truncated Intron 1 that also allows for robust expression of RFK-1B in vegetative tissue will aid future efforts to determine how RFK-1B acts as a poison. For example, full length Intron 1 contains 395 nucleotides, and its repetitive sequences make it difficult to construct with synthetic DNA technology. In contrast, the truncated version of Intron 1 examined in this study contains only 105 nucleotides and lacks repeats, which will simplify future efforts to construct synthetic *rfk-1* variants.

In this study we also constructed a plasmid-based transformation vector that may be of use to researchers studying Neurospora fungi. The vector, pNR304.1, is useful for placing transgenes under control of the *tcu-1* promoter. One caveat with respect to the use of this vector is that it deletes the *tcu-1* coding region, resulting in a *tcu-1* knockout strain. While this may be undesirable for some studies, we did not detect phenotypic differences between *tcu-1^+^*and *tcu-1Δ* genotypes in our radial or linear growth assays (Figure 2), suggesting that deletion of *tcu-1* might have little impact on *N. crassa* physiology when grown under standard laboratory conditions.

The discovery of RFK-1B, along with the finding that the poison and antidote system of *Sk-2* can be phenotypically expressed in vegetative tissue, should greatly facilitate future efforts to elucidate the molecular processes used by *Sk-2* to achieve biased transmission to the next generation through spore killing. *Sk-2* was discovered along with two other Neurospora Spore killers over 40 years ago, the single gene element *Sk-1*, and the complex element *Sk-3* (Turner and Perkins 1979; Svedberg et al. 2021). In many ways, *Sk-2* and *Sk-3* are highly similar. They both span millions of base pairs of Chromosome III, they are both associated with recombination blocks enforced by several major inversions, and they both use RSK variants to resist spore killing (Campbell and Turner 1987; Hammond et al. 2012; Svedberg et al. 2018). However, their inversions appear to have evolved independently, and efforts to identify an *rfk-1-*like gene that is required for spore killing by *Sk-3* have been unsuccessful (Svedberg et al. 2021; Velazquez et al. 2022). Thus, the evolutionary history of *Sk-2* and *Sk-3* is both intriguing and perplexing.

Elucidating how RFK-1B and RSK^Sk-2^ function may provide insight into the identity of the *Sk-3* poison and help explain the unusual relationship between these two complex selfish genetic elements.

## ACKNOWLEDGEMENTS

This study was made possible by the pioneering work of Barbara Turner, Namboori Raju, David Perkins, and others. We thank members of the Hammond Laboratory for technical assistance with this work. We also thank the Fungal Genetics Stock Center for FGSC 10340 and for serving as a repository for the Spore killer strains (McCluskey et al. 2010). TH conceived the project, designed the experiments, provided technical assistance as needed, analyzed the results, and wrote the manuscript. SM constructed plasmids and strains with pSM, CSM, TSM, and RSM names. NR constructed plasmids and strains with pNR, CNR, TNR, and RNR names. SM performed the experiments depicted in Figures 2, 3, and 4. NR performed the experiments depicted in Figure 1E and 1F, as well as replicate assays to the one presented in Figure 2, and preliminary experiments on the role of Intron 1 in *rfk-1^+^* expression. NR edited the manuscript. All authors reviewed and approved the final version of the manuscript. This work was supported by awards to TMH from the National Science Foundation (1615626 / 2005295).

