## Supplementary material for "RNA editing controls gene drive by a Neurospora Spore killer": Figure S1

Synthetic DNA 1 (*AH36/rfk-1*$^{+,Int1\Delta}$)

TTTT**GCGGCCGC**ATTGATACCGAGTCTTTCCGTTCTTAAGGTTGGAGTGAGGATATGATCCGGCACGTC
GAAGGAGGAACTATGGTAAATAAGTCAGTCATCATCAAGGAAAAGAACAGTATTAGGTGCCTTCAAGTT
ACCTACCTTCTCCGACGGGGGCCCTGCCTTCCCTTAGTTCTCTCATGAAGCTGGAAGTTATATCCTAAC
CCACTACTGACAAAACAAAGCAAGAGCTGCTGCCCCAACTATAGACAGCACGCTTTTCCACCTCAGTTG
GGGCACCTAGAAAGCTATAAGATCCCTCTTCCCCCGGCCCAACCTCTCCTCAGAATTTCTTTTTTTCTC
CAACATTGTTAAGAACTTTGTTTTTGGAAAATGGCCTGCCCCACAGGGTTTTTTACCGCTCTTTTTGGC
AAACTCCTCACCATCCCCATTTGGGTGTTGGTGTTTGTATTCAATGCTCTGTTTGTCTTCCCCCGGTTT
TGGGTCTGGCGCTTACCACCACGGCTGCAACACGAGCAGCAGCAGCAGGACCGGAACGATGACGAATGG
CAGCGGCAGCAACAGGACAGGCGGGTTGTGGTTTGGCACCCACCGCCCCCTCCAGACGTGGAGATGGCC
CTCCAAGACAATCCCACTGCCGCCCCCGCAGAGCCGGCTGACCTCGACCACCCAGCGCCGTAGTGGCGG
CGCTGGGTGGCCGACGAGTAGGTCAATGCTATTCCCAGATTATGAAATGTATCGCTGACAGTTGCACAC
CAGTGCCTACCCGGCCGTCCACTTCTGCGTGACCGCAGCCAATGCGGTCACGCAGGGGTTGTAATTCCA
CGTGAGCATTCCCCACCTTCTCTCGGGACCGACTTCCGTATCAACCCCAAATTTATCGGACTGACCCGT
CCGAATCAAGGCGAACCGAGAGGACACAGACAAGGCCCACGTCCGCCATCAGCATTCCCAGCTGGCCGA
CCGCACCGCCGCAACTCCCACTTTACCTCAACACCAGAATACGGAATCGGTACATCGACAGCAGCATCA
TCATCATCAATATCACCACCTCCACTTGGCGCGCACTTGCGGAAAACGTCCCGCTACACCGT**GCGGCCG**
**C**TTTT

**Figure S1** Synthetic DNA 1. The sequence of Synthetic DNA 1 is shown. Synthetic DNA 1 contains *AH36/rfk-1*$^{Int1\Delta}$. The start codon, early stop codon, and late stop codon are underlined. Two *Not*I recognition sequences (bold font) were included for cloning purposes. Intron 2 and Intron 3 are indicated with gray highlighting.
