## Supplementary material for "RNA editing controls gene drive by a Neurospora Spore killer": Figure S2

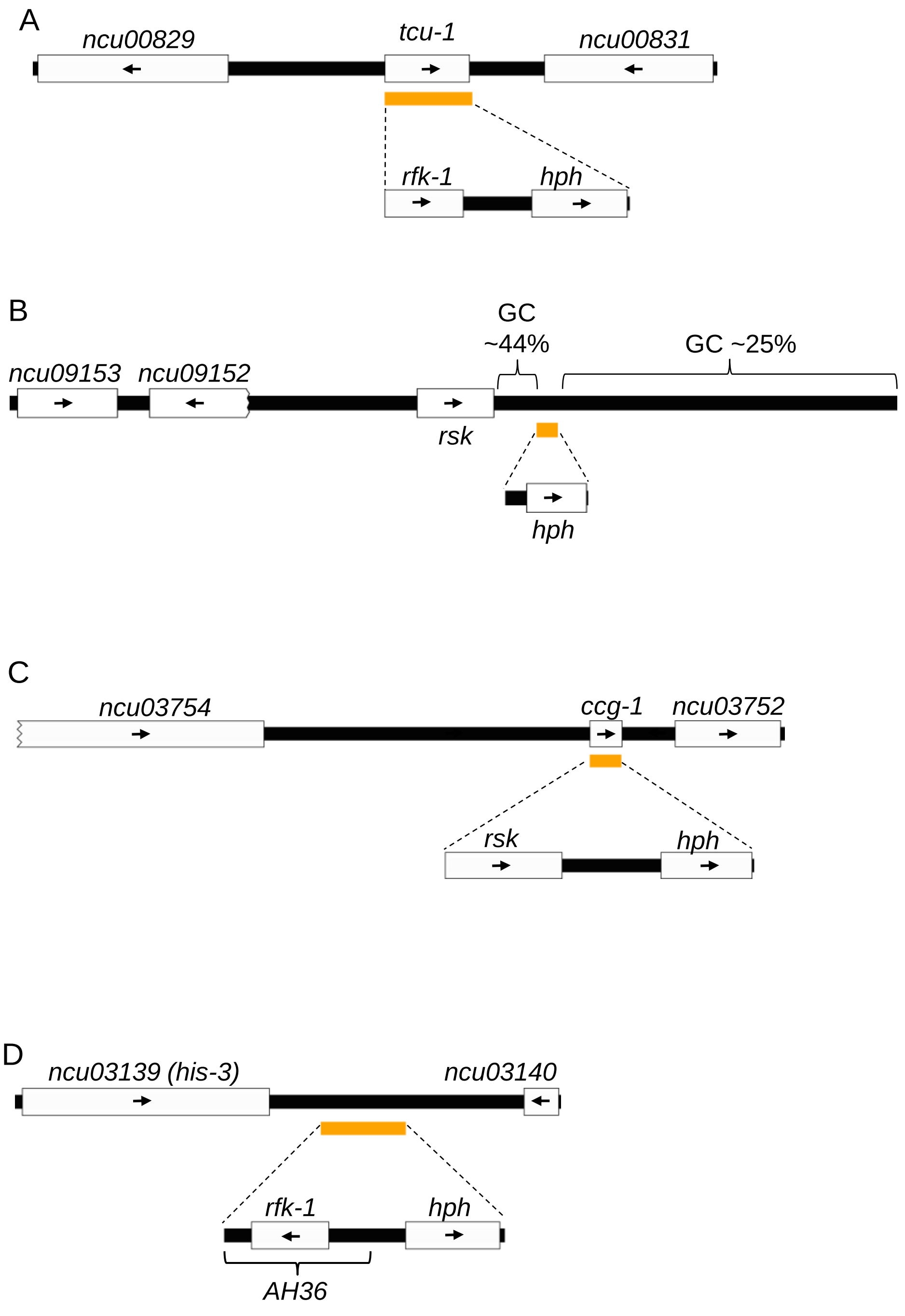
A
ncu00829
tcu-1
ncu00831
rfk-1
hph
B
GC
~44%
GC ~25%
ncu09153
ncu09152
rsk
hph
C
ncu03754
ccg-1
ncu03752
rsk
hph
D
ncu03139 (his-3)
ncu03140
rfk-1
hph
AH36

**Figure S2** Transgene integration maps. (A) The *N. crassa tcu-1* locus on Chromosome I is shown in the diagram. The relative positions of *N. crassa* genes *ncu00829*, *tcu-1*, and *ncu00831* are indicated. Coding regions are depicted as white rectangles. Intron positions are not depicted. The directions of transcription are indicated with black arrows. The diagram was drawn to scale and the orange bar represents the 936 bp of DNA that were deleted and replaced with *rfk-1* sequences and a selectable marker (*hph*$^+$) in ISU-4956 and ISU-4957. (B) The relative positions of *N. crassa* genes *ncu09153*, *ncu09152*, and *rsk* on Chromosome III in *Sk-2* strain FGSC 7426 (Svedberg et al. 2018) are shown. A putative start codon for *ncu09152* was not identified (indicated by the jagged edge of the *ncu09152* white rectangle). The diagram was drawn to scale with the orange bar representing the 362 bp of DNA that were deleted and replaced with a selectable marker (*hph*$^+$) to create ISU-4958. Most of the approximately 6800 bp of DNA sequence depicted to the right of *rsk* in the diagram has a low GC content (no genes detected). (C) The relative positions of *N. crassa* genes *ncu03754*, *ccg-1*, and *ncu03752* on Chromosome V are indicated. Only part of the *ncu0375*4 coding sequence is represented in the diagram. The diagram was drawn to scale with the orange bar representing the 354 bp of DNA that were deleted and replaced with *rsk*$^{Sk\text{-}2}$ coding sequences and a selectable marker (*hph*$^+$) to create the *ccg-1*$^{\Delta}$*::rsk*$^{Sk\text{-}2}$ allele in ISU-4959 and its descendants (ISU-4961, ISU-4962, and ISU-4963). (D) The *N. crassa his-3* locus on Chromosome I is depicted in the diagram. The position of the *his-3*$^+$ coding sequence is indicated with a white rectangle, as is the position of the coding sequence for gene *ncu03140*. Intron positions are not indicated. The diagram was drawn to scale and the orange bar represents the 919 bp of intergenic sequence that was deleted and replaced with *AH36*/*rfk-1* sequences and a selectable marker (*hph*$^+$) in ISU-4954 and ISU-4955.
