## Supplementary material for "RNA editing controls gene drive by a Neurospora Spore killer": Figure S3

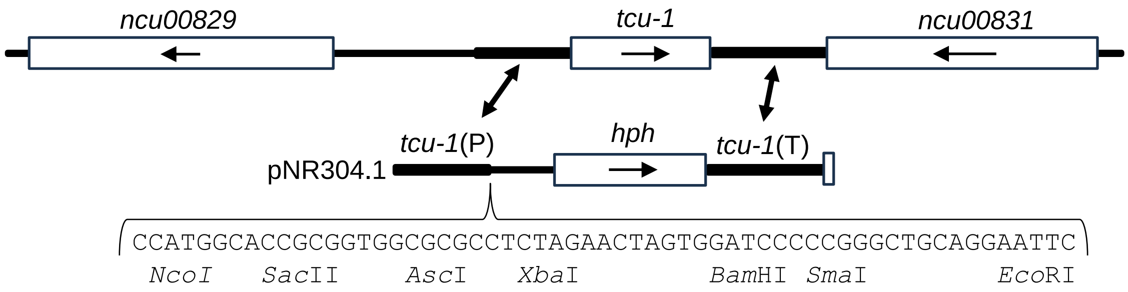


**Figure S3** Transformation vector pNR304.1. A map of the *tcu-1* locus and the *tcu-1* replacement vector, pNR304.1 (GenBank PX833383), is shown. The multiple cloning site contains seven unique restriction endonuclease recognition sites that can be used to insert coding sequences along with a 3′UTR/Terminator. The ATG of the *Nco*I site is intended to serve as the start codon for the inserted coding sequence. The vector is designed to replace the entire *tcu-1* coding region, and two nucleotides of the *tcu-1* 3'UTR, of a transformation host, with a desired coding sequence, 3′UTR/Terminator, and *hph*$^{+}$ by double homologous recombination. The recombination sites are indicated with double sided arrows. The diagram was drawn to scale, with the *hph*$^{+}$ coding region (rectangle) representing 1026 bp.
